# Drift barriers to quality control when genes are expressed at different levels

**DOI:** 10.1101/058453

**Authors:** K Xiong, JP McEntee, DJ Porfirio, J Masel

## Abstract

Gene expression is imperfect, sometimes leading to toxic products. Solutions take two forms: globally reducing error rates, or ensuring that the consequences of erroneous expression are relatively harmless. The latter is optimal, but because it must evolve independently at so many loci, it is subject to a stringent “drift barrier” – a limit to how weak the effects of a deleterious mutation s can be, while still being effectively purged by selection, expressed in terms of the population size *N* of an idealized population such that purging requires *s* < −1/*N*. In previous work, only large populations evolved the optimal local solution, small populations instead evolved globally low error rates, and intermediate populations were bistable, with either solution possible. Here we take into consideration the fact that the effectiveness of purging varies among loci, because of variation in gene expression level and variation in the intrinsic vulnerabilities of different gene products to error. The previously found dichotomy between the two kinds of solution breaks down, replaced by a gradual transition as a function of population size. In the extreme case of a small enough population, selection fails to maintain even the global solution against deleterious mutations, explaining the non-monotonic relationship between effective population size and transcriptional error rate that was recently observed in experiments on *E. coli*, *C. elegans* and *Buchnera aphidicola*.

## INTRODUCTION

In classical population genetic models of idealized populations, the probability of fixation of a new mutant depends sharply on the product of the selection coefficient s and the population size *N*. As s falls below −1/*N*, fixation probabilities drop exponentially, corresponding to efficient selective purging of deleterious mutations. For *s* > −1/*N*, random genetic drift makes the fate of new mutants less certain. This nonlinear dependence of fixation probability on *sN* has given rise to the “drift barrier” hypothesis (Lynch 2007), which holds that populations are characterized by a threshold or “barrier” value of the selection coefficient s, corresponding to the tipping point at which the removal of deleterious mutations switches between effective and ineffective. In idealized populations described by Wright-Fisher or Moran models, the drift barrier is positioned at *s* = ~−1/*N*. Drift barriers also exist, albeit sometimes with less abrupt threshold behavior, in more complex models of evolution in which some assumptions of an idealized population are relaxed (Good and Desai 2014).

The drift barrier theory argues that variation among species in their characteristic threshold values for *s*, thresholds that are equal by definition to the inverse of the selection effective population size *N*_*e*_, can explain why different species have different characteristics, e.g. streamlined versus bloated genomes (Lynch 2007). The simplest interpretation of the drift barrier would seem to imply that large-*N*_*e*_ species show stricter quality control over all biological processes, e.g. higher fidelity in DNA replication, transcription, and translation, than small-*N*_*e*_ species, because molecular defects in quality control mechanisms are less effectively purged in the latter (Lynch 2010; Traverse and Ochman 2016a).

However, the data reveals more complex patterns. Unsurprisingly, *Buchnera aphidicola*, which has exceptionally low *Ne* (Mira and Moran 2002; Rispe *et al.* 2004), has a higher transcriptional error rate, at 4.67×10^−5^ (Traverse and Ochman 2016b), than the error rate 4.1×10^−6^ previously reported for *Caenorhabditis elegans* (Gout *et al.* 2013). But to the surprise of the authors, the error rate in large-*N*_*e*_ *Escherichia coli* is highest of all, at 8.23×10^−5^ (Traverse and Ochman 2016b).

A more refined drift barrier theory can explain these findings. As the fitness burden accumulates from the slightly deleterious mutations that a small-*N*_*e*_ species cannot purge, some forms of quality control may evolve as a second line of defense. The ideal solution is to purge all deleterious mutations, even those of tiny effect; when this first line of defense fails, the second line of defense is to ameliorate the cumulative phenotypic consequences of the deleterious mutations that have accumulated (Frank 2007; Rajon and Masel 2011; Warnecke and Hurst 2011; Lynch 2012; Wu and Hurst 2015). In some circumstances, as described further below, strict quality control can act as such an amelioration strategy (Rajon and Masel 2011). The existence of two distinct lines of defense complicates the naive drift barrier logic that large-*N*_*e*_ species should generally exhibit stricter quality control in all molecular processes. The superior performance of large-*N*_*e*_ species in a primary line of defense other than quality control may remove any advantage of strict and costly quality control as a secondary line of defense. This creates a seemingly counter-intuitive pattern in quality control, in which small-*N*_*e*_ species can evolve more faithful processes than large-*N*_*e*_ species such as *E. coli*.

The existence of two substantively different lines of defense was first proposed by Krakauer and Plotkin (2002), who contrasted the “redundancy” of robustness to the consequences of mutational errors with the “antiredundancy” of hypersensitivity to mutations. By positing that redundancy had a cost, they showed that the superior cost-free solution of antiredundancy was available only with large *N*_*e*_, giving small-*N*_*e*_ species higher levels of “redundancy”.

A related argument was made by Rajon and Masel (2011) in the context of mitigating the harms threatened by errors in molecular processes such as translation. Rajon and Masel (2011) distinguished between “local” solutions, where a separate solution is required at each locus, and “global” solutions that can deal with problems at many loci simultaneously. The evolution of extensive quality control mechanisms was deemed a global solution because a single mutation impacting general quality control mechanisms can affect the prevention of gene expression errors at many loci. Note that quality control includes not only mechanisms such as proofreading for preventing errors from happening in the first place, but also mechanisms that reduce downstream damage from errors, e.g. degradation of mRNA molecules that seem faulty. Global quality control should come with a cost in time or energy. The alternative, local solution is to have a benign rather than a strongly deleterious “cryptic genetic sequence” at each locus at which expression errors might occur, making the consequence of an error at that locus relatively harmless. In contrast to the global solution, these local solutions bear no direct fitness cost, but because selection at any one locus is weak, mutations at any one locus pass more easily through the drift barrier, making them more difficult to maintain than global solutions.

Both the quality control of Rajon and Masel (2011) and the “redundancy” of Krakauer and Plotkin (2002) to the consequences of mutations are global across loci, and also costly (the former costly by design, the latter costly as a consequence of scaling decisions). Meantime, both the “local” solutions of Rajon and Masel (2011) and the “antiredundancy” of Krakauer and Plotkin (2002) carry no true fitness cost but instead require a large-*N*_*e*_ drift barrier and/or face a “cost of selection” (Haldane 1957) as limits to their adaptation. A mutation disrupting a solution specific to a single locus requires a large value of *N*_*e*_ for its purging, whereas a mutation disrupting a global quality control mechanism will have large fitness consequences and so be easier to purge. The higher-fitness solution is the local one, but it is evolutionarily achievable only with large *N*_*e*_. With small *N*_*e*_, we instead expect global solutions such as extensive (and costly) quality control.

Selection to achieve the local solution by purging deleterious mutations to cryptic sequences (leaving in place genotypes whose cryptic genetic sequences are benign) may be difficult and hence restricted to high-*N*_*e*_ populations. There are, however, reasons to believe that it is not impossible. For example, when the error in question is reading through a stop codon, the local cryptic genetic sequence is the 3′UTR, which is read by the ribosome. One option for a more benign form of this cryptic sequence is the presence of a “backup” stop codon that provides the ribosome with a second and relatively early chance to terminate translation. Such backup stops are common at the first position past the stop in prokaryotes (Nichols 1970). In *Saccharomyces cerevisiae*, there is also an abundance of stop codons at the third codon position past the stop (Williams *et al.* 2004). Moreover, conservation at this position depends strongly on whether or not the codon is a stop, and the overrepresentation of stops at this position is greater in more highly expressed genes (Liang *et al.* 2005). In some ciliates, where the genetic code has been reassigned so that UAA and UAG correspond to glutamine, this overrepresentation is much more pronounced (Adachi and Cavalcanti 2009). As with the consequences of erroneous readthrough, selective pressure on erroneous amino acid misincorporation and/or misfolding (Drummond and Wilke 2008), and on erroneous protein-protein interactions (Brettner and Masel 2012) are also strong enough to shape protein expression and interaction patterns. In the case of transcriptional errors, while both *E. coli* and *B. aphidicola* have high error rates, only *E. coli* shows signs of having evolved a first line of defense in the form of a decreased frequency with which observed transcriptional errors translate into non-synonymous changes, relative to randomly sampled transcriptional errors (Traverse and Ochman 2016a).

Rajon and Masel (2011) found that for intermediate values of *N_*e*_* that correspond strikingly well to many multicellular species of interest, the evolutionary dynamics of the system were bistable, with either the global or the local solution possible. This is a natural consequence of a positive feedback loop; in the presence of a strict global quality control mechanism, specialized solutions at particular loci are unnecessary and mutations destroying them pass through the drift barrier (we use the expression “pass through the drift barrier” to mean that 0 > *s* > −1/*N*), with their subsequent absence increasing the demand for quality control. Similarly, when specialized solutions predominate, the advantage to quality control is lessened, and resulting higher error rates further increase selection for many locally specialized solutions. If true, this bistability suggests that historical contingency, rather than current values of *N*_*e*_, determine which processes are error-prone vs. high-fidelity for populations at intermediate *N*_*e*_.

In the current work, we note that the model of Rajon and Masel (2011) contained an unrealistic symmetry, namely that the fitness consequence of a molecular error at one locus was exactly equal to that at any other loci. Here we find that with reasonable amounts of variation among loci (e.g. in their expression level or the per-molecule damage from their misfolded form), the bistability disappears. Intermediate solutions evolve instead, where cryptic deleterious sequences are purged only in more highly expressed genes, and quality control evolves to intermediate levels. Variation among loci does not change the previous finding that evolvability tracks the proportion of loci that contain a benign rather than a deleterious cryptic sequence.

The high rate of transcriptional error in *B. aphidicola* can be explained by adding a second bias toward deleterious mutations (in error rate), and hence a second drift barrier to our model. *B. aphidicola* and *E. coli* have high error rates for different reasons; high-fidelity quality control is redundant and unnecessarily expensive in *E. coli*, but unattainable in *B. aphidicola*, leading to similarly high transcriptional error rates.

## METHODS

In the following sections, we describe the computational model used to simulate the evolution of different solutions to errors in gene expression. All simulations were run with Matlab (R2014a). Source code for the simulations is available at https://github.com/MaselLab/.

### Fitness

We follow the additive model of Rajon and Masel (2011), as outlined below, with a few important modifications to accommodate variation in gene expression levels. The model's canonical example is the risk that a ribosome reads through a stop codon during translation.

The global mitigation strategy is to improve quality control of this gene expression subprocess. We assume that additional quality control that reduces the error rate ρ by some proportion consumes a certain amount of time or comparable resource. Relative to a generation time of 1 in the absence of quality control costs, this gives generation time 1 + *δ* ln(1/*ρ*), where *δ* scales the amount of resources that could have been used in reproduction but are redistributed to quality control. Malthusian fitness is the inverse of generation time, giving

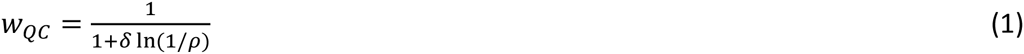

Following Rajon and Masel (2011), we set *δ* = 10^−2.5^, such that reducing *ρ* from 10^−2^ to 10^−3^ corresponds to a 0.7% reduction in fitness.

When a readthrough error happens, with frequency *ρ*, the consequences for fitness depend on the nature of the “cryptic sequence” that lies beyond the stop codon in the 3′UTR. The consequences of mistakes, mutational or otherwise, have a bimodal distribution, being either strongly deleterious (often lethal), or relatively benign, but rarely in between (Eyre-Walker and Keightley 2007; Fudala and Korona 2009). For example, a strongly deleterious variant of a protein might misfold in a dangerous manner, while a benign variant might fold correctly, although with reduced activity. We assume that alternative alleles of “cryptic genetic sequences” can be categorized according to a benign/deleterious dichotomy.

The local mitigation strategy, the alternative to global quality control, is thus for each cryptic sequence to evolve away from “deleterious” options and toward “benign” options. The local strategy of benign cryptic sequences has no direct fitness cost, but it is nevertheless difficult to evolve at so many loci at once. In contrast, expressing deleterious cryptic sequences has an appreciable cost. This cost scales both with the base rate of expression of the gene, and the proportion *ρ* of gene products that include the cryptic sequence.

Let the expression of gene *i* be *E*_*i*_. We assign the concentration *E*_*i*_ of protein molecules of type *i* by sampling values of *E*_*i*_ from a log2-normal distribution with standard deviation *σ*_*E*_. We define *D* to be the total frequency of protein expression that would be highly deleterious if expressed in error:

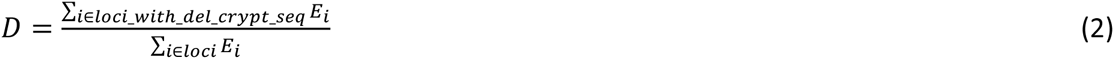

where the numerator sums only over loci that are deleterious and the denominator sums over all loci. This normalization cancels out the effect of the mean value of *E*_*i*_. We assume the costs of deleterious readthrough are additive across genes, based on the concept that misfolded proteins (Thomas *et al.* 1995) may aggregate in a non-specific and harmful manner with other proteins and/or membranes (Kourie and Henry 2002), or may simply be expensive to dispose of (Goldberg 2003). After the stop codon is read through, translation will usually end at a backup stop codon within the 3′UTR. Under the assumption of additivity, readthrough events will reduce fitness by *cρD*, where *c* represents the strength of selection against misfolded proteins. Geiler-Samerotte et al. (2011) found that an increase in misfolded proteins of approximately 0.1% of total cellular protein molecules per cell imposed a cost of about 2% to relative growth rate. This gives an estimate of *c* = 0.02/0.1% = 20.

Readthrough involving benign cryptic sequences does not incur this cost. However, when all cryptic sequences are benign (i.e. *D* = 0), nothing stops *ρ* from increasing to unreasonably large values, i.e. *ρ* > 0.5, which makes “erroneous” expression into the majority (and hence the “new normal") form. To avoid this scenario, we add a cost in fitness *cρ*^2^(1-*D*), whose impact is felt only at high values of *ρ*. One possible biological interpretation of this second order term is that with probability *ρ*^2^, readthrough occurs not just through the regular stop codon, but also through the backup stop codon at the end of the benign cryptic genetic sequence. To reflect the effects of the double-error scenario under this interpretation, we therefore multiplied the second order term by the probability *μ*_*del*_/(*μ*_*del*_+*μ*_*ben*_) that a neutrally evolving cryptic sequence will be deleterious, where *μ*_*del*_ is the rate of deleterious-to-benign mutations and *μ*_*ben*_ the reverse rate. Other double-error interpretations might involve different constants. In our case, the fitness component representing the cost of misfolded proteins is given by

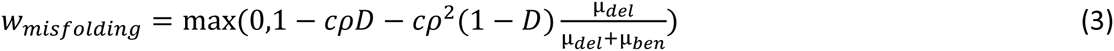

Eq. 3 is a natural extension of the additive model of Rajon and Masel (2011), generalizing to the case of variation in the degree of importance of cryptic loci. Where previous work referred to the number *L*_*del*_ of loci having the deleterious rather than benign form, we now distinguish between two measures, *L*_*del*_ and *D*, the latter reporting the proportion of gene product molecules rather than gene loci.

Rajon and Masel (2011) also obtained near-identical results using a very different, multiplicative model. While this suggests that the exact function form of Eq. 3 is unimportant, we chose the additive Eq. 3 model as the more reasonable of the two options. The multiplicative model is premised on loss-of-function of the wild-type proteins, which likely has negligible impact for small losses of a protein whose activity is already close to saturation. In contrast, the additive model is premised on gain-of-negative-function effects of misfolded proteins. These plausibly constitute a major burden on fitness, through a combination of toxicity, disposal costs, and resources spent to replace a faulty molecule with a normal one.

To study evolvability, let a subset of *K* (typically 50) out of the *L* (typically 600 or more) loci affect a quantitative trait *x*, selection on which creates a third fitness component. Error-free expression of locus k, occurring with frequency 1-*ρ*, has quantitative effect *αk*, while expression that involves a benign version of the cryptic sequence has quantitative effect *α*_*k*_ + *β*_*k*_. Expression that involves a deleterious version of the cryptic sequence is assumed to result in a misfolded protein that has no effect on the quantitative trait. We assume that expression level *E*_*k*_ is constant and already factored into values of *α*_*k*_ and *β*_*k*_. This gives

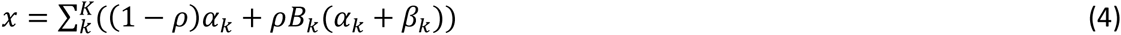

where *B*_*k*_ = 1 indicates a benign cryptic sequence and *B* = 0 a deleterious one. As in Rajon and Masel (2011), we impose Gaussian selection on *x* relative to an optimal value *x*_*opt*_

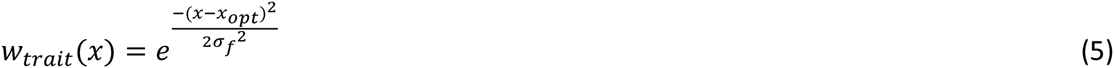

where *σ*_*f*_ = 0.5.

Putting the three fitness components together, the relative fitness of a genotype is given by the product

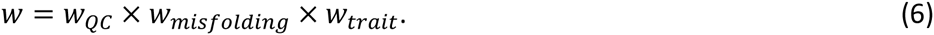

### Variance in expression levels

We estimated the variance in expression 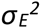 from PaxDB (Wang *et al.* 2012; Wang *et al.* 2015), which is based on data released by the Global Proteome Machines (GMP) and other sources. We inferred *σ*_*E*_ equal to 2.24 (based on GMP 2012 release) or 3.31 (GMP 2014 release), for *S. cerevisiae*, and 2.93 (GMP 2014 release) for *S. pombe*. Note that while our quantitative estimate of *σ*_*E*_ comes from variation in the expression levels of different proteins, consideration of variation along other lines might make a standard deviation of 2.25 into a conservative underestimate of the extent of variation. See Fig. S2 for an exploration of this parameter value.

### Mutation

There are six kinds of mutation: 1) conversion of a deleterious cryptic sequence to a benign form, 2) conversion from benign to deleterious, 3) change to the error rate *ρ,* 4) change in the *α* value of one of the *K* quantitative trait genes, 5) change in the *β* value of one of those *K* genes, and 6) the co-option of a cryptic sequence to become constitutive, replacing the value of replacing *αk* with that of *α*_*k*_+*β*_*k*_ and re-initializing *B*_*k*_ and *β*_*k*_.

It is this sixth kind of mutation that is responsible for the evolvability advantage of the local solution of benign cryptic sequences, providing more mutational raw material by which *x* might approach *x*_*opt*_ (Rajon and Masel 2011; Rajon and Masel 2013). The mutational co-option of a deleterious sequence (*B* = 0) is too strongly deleterious to be favored, even when replacing *α*_*k*_ and *βk* might be advantageous. In other words, only benign cryptic sequences are available for mutational co-option. We use the term co-option of a 3′UTR readthrough sequence to refer to the case when a stop codon is lost by mutation, and not just read through by the ribosome (Giacomelli *et al.* 2007; Vakhrusheva *et al.* 2011; Andreatta *et al.* 2015). Mutational co-option for mimicking the consequences of errors other than stop codon readthrough might involve mutations that change expression timing to make a rare protein-protein interaction common, or switch a protein's affinity preference between two alternative partners.

Because we use an origin-fixation approach to simulate evolution (see below), only relative and not absolute mutation rates matter for our outcomes, with the absolute rates setting only the timescale – our rates are therefore effectively unitless. We use the same mutation rates as Rajon and Masel (2011), reduced ten-fold for convenience. Each locus with a benign cryptic sequence mutates to deleterious at rate *μ*_*del*_ = 2.4×10^−8^, while deleterious loci mutate to benign less often, at rate *μ*_*ben*_ = 6×10^−9^. Changes to the error rate *ρ* occur at rate *μ*_*ρ*_ = 10^−6^, while the *α* and *β* values of quantitative loci each change with rates *μ*_*α*_ = 3×10^−7^ and *μ*_*β*_ = 3×10^−8^, respectively. Mutational co-option occurs at each quantitative locus at rate *μ*_*coopt*_ =2.56×10^−9^.

Each mutation to *ρ* increases log_10*ρ*_ by an amount sampled from Normal(*ρ*_*bias*_,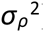). By default, we set *ρ*_*bias*_ = 0 and *σ*_*ρ*_ = 0.2. To study extremely small populations with drift barriers to evolving even a global solution, we set *ρ*_*bias*_ = 0.256 and 0.465, corresponding to ratios of *ρ*-increasing mutations: *ρ*-decreasing mutations of 9:1 and 99:1, respectively.

A similar scheme for α and *β* might create, in the global solution case of relaxed selection, a probability distribution of *β* whose variance increases in an unbounded manner over time (Lande 1975; Lynch and Gabriel 1983). Following previous work (Rajon and Masel 2011; Rajon and Masel 2013), we therefore let mutations alter *α* and *β* by an increment drawn from a normal distribution with mean −*α/a* or −*β/a,* with *a* set to 750, and with standard deviation of *σ*_*m*_/*K* in both cases, with *σ*_*m*_ set to 0.5. In the case of neutrality, this mutational process eventually reaches a stationary distribution with mean 0 and standard deviation as calculated in Eq. S3 of Rajon and Masel (2011):

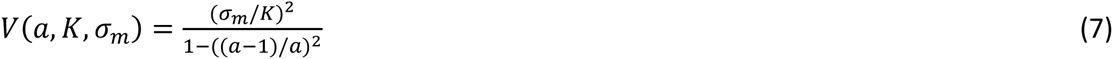

A co-option at gene *k* changes the gene's quantitative effect to

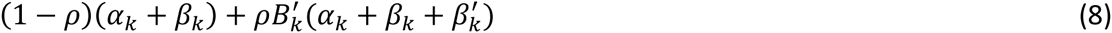

where 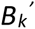 and 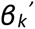 are the state and the quantitative effect of a new cryptic sequence created by co-option. Following a co-option mutation at locus *k*, we set the new *B*_*k*_ equal to 1 or 0 with probabilities proportional to *μ*_*ben*_ and *μ*_*del*_, and resample the value of *β*_*k*_ from Normal(0, *V*(*a*, *K*, *σ*_*m*_)).

### Evolutionary simulations by origin-fixation

We model evolution using an approach known as “weak mutation” (Gillespie 1983), or “origin-fixation” (McCandlish and Stoltzfus 2014). This approximation of population genetics is accurate in the limit where the waiting time until the appearance of the next mutation destined to fix is substantially longer than its subsequent fixation time. The population can then be approximated as genetically homogeneous in any moment in time. While unrealistic for higher mutation rates and larger population sizes, origin-fixation models are computationally convenient. Still more importantly, origin-fixation models, unlike more realistic models with segregating variation, allow the location of the drift barrier to be set externally in the form of the value of the parameter *N*, rather than having the location of the drift barrier emerge from complicated linkage phenomena within the model. Fortunately, for quantitative traits affected by multiple cryptic loci, most evolvability arises from diversity of the effects of co-option of different loci, rather than among the diversity of the effects of co-option from different starting genotypes (Rajon and Masel 2013). This allows us to study evolvability (in the population sense of Wagner (2008)) even in the absence of genetic diversity that is imposed by the origin-fixation formulation.

Our computationally efficient implementation of origin-fixation dynamics is described in detail in the Supplement, simulating a series of mutations that successfully fix, and the waiting times between each.

### Initialization and convergence

We initialized the trait optimum at *x*_*opt*_ = 0. We could have initialized all values of *α*_*k*_ and *β*_*k*_ at zero. However, at steady state, variation in 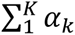 and 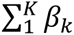 is far lower than would be expected from variation in *α*_*k*_ and *β*_*k*_ – this emerges through a process of compensatory evolution (Rajon and Masel 2013). Allowing a realistic steady state to emerge in this way is computationally slow under origin-fixation dynamics, especially when *N* is large. We instead sampled the initial values of *α*_*k*_ and *β*_*k*_ from Normal(0, *V*(*a*, *K*, *σ*_*m*_)), where *V*(*a*, *K*, *σ*_*m*_) is defined by Eq. 7, and then subtracted 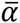 from *α*_*k*_ and 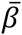 from *β*_*k*_, where 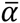 and 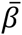 are the means of a genotype across each of its quantitative loci *k.* This process initializes *α*_*k*_ and *β*_*k*_ to have variances equal to those of the stationary distributions, while the overall trait value is initialized at the optimal value, zero. This procedure greatly reduces the burn-in computation time needed to achieve a somewhat subtle state of negative within-genotype among-loci correlations. We confirmed that subsequent convergence of the variance of 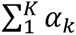 was fast, occurring in less than 1000 steps, where a “step” is defined to be the fixation of one mutation. We expect log_10_*ρ*, *D*, and variation in *β*_*k*_, to converge even faster than variation in *α*_*k*_.

For the low-*ρ* initial conditions, *ρ* was initialized at 10^−5^, and we initialized the benign vs. deleterious status of cryptic sequences at the neutral mutational equilibrium, choosing exactly *L*×*μ*_*del*_/(*μ*_*del*_+*μ*_*ben*_) (rounded to the nearest integer) to be deleterious, independently of their different values of *E*. For the high-*ρ* initial conditions, we set *ρ* to 10^−1^, and made all cryptic sequences benign.

We ran simulations for 10^5^ steps, recording information at fixed times (measured in terms of waiting times), corresponding to approximately every 1000 steps on average, and hence yielding about 100 timepoints. To summarize the evolutionary outcome, we calculated the arithmetic means of log_10_*ρ*, of *L*_*del*_, and of *D* among the last 20 timepoints, i.e. approximating steps 0.8×10^5^−1×10^5^.

### Evolvability

After adaptation to a trait optimum of *x*_*opt*_ = 0 had run to convergence (i.e. after 10^5^ steps), we changed *x*_*opt*_ to 2, forcing the quantitative trait to evolve rapidly. This allows the co-option of benign cryptic sequences an opportunity to increase evolvability. We measured evolvability in two ways: as the inverse of the waiting time before trait *x* exceeded 1, and the inverse of the waiting time before the population recovered half of the fitness it lost after *x*_*opt*_ changed. By default, we present results showing evolvability as time to fitness recovery; evolvability as time to trait recovery is shown only in Fig. S3.

We want our measures of evolvability to reflect a genotype's potential to generate beneficial mutations, but this goal was complicated by population size. A large population finds a given beneficial mutation faster than a small population does, inflating the total fixation flux 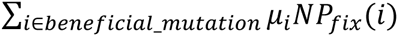, where *μ*_*i*_*N* is the influx of mutations of beneficial type *i* and *P*_*fix*_ is their probability of fixation (the latter described by Eq. 9 in the supplemental material), in direct proportion to population size. We therefore divided our evolvability measures by the population size to correct for this effect. This normalization converts the population-level evolvability measure into a measure of the population-size-independent evolvability of a single individual that has the genotype of interest.

## RESULTS

Recall that in the absence of variation in expression among genes, there are two solutions to handle erroneous expression due to stop codon readthrough: at high population size *N*, the local solution purges all deleterious cryptic sequences, making high rates of readthrough harmless, while at low *N*, the global solution reduces the rate of readthrough, allowing deleterious cryptic sequences to accumulate near-neutrally. At intermediate *N*, we see bistability, with either solution possible, depending on starting conditions (Fig. 1, *σ* _*E*_ = 0). It is important to note that we use the word “bistability” loosely. Strictly speaking, bistability means that the system has two stable steady states (here a state is defined by readthrough rate and the exact property of each cryptic sequence), i.e. two attractors. But in a stochastic model, there are no attractors in the strict sense of the word, only a stationary distribution of states. We use the term bistability to refer to the case where the stationary distributions of states has two modes. Transitions between the two modes are rare, therefore the two modes can be loosely interpreted as the two attractors of the system.

**Figure 1:**
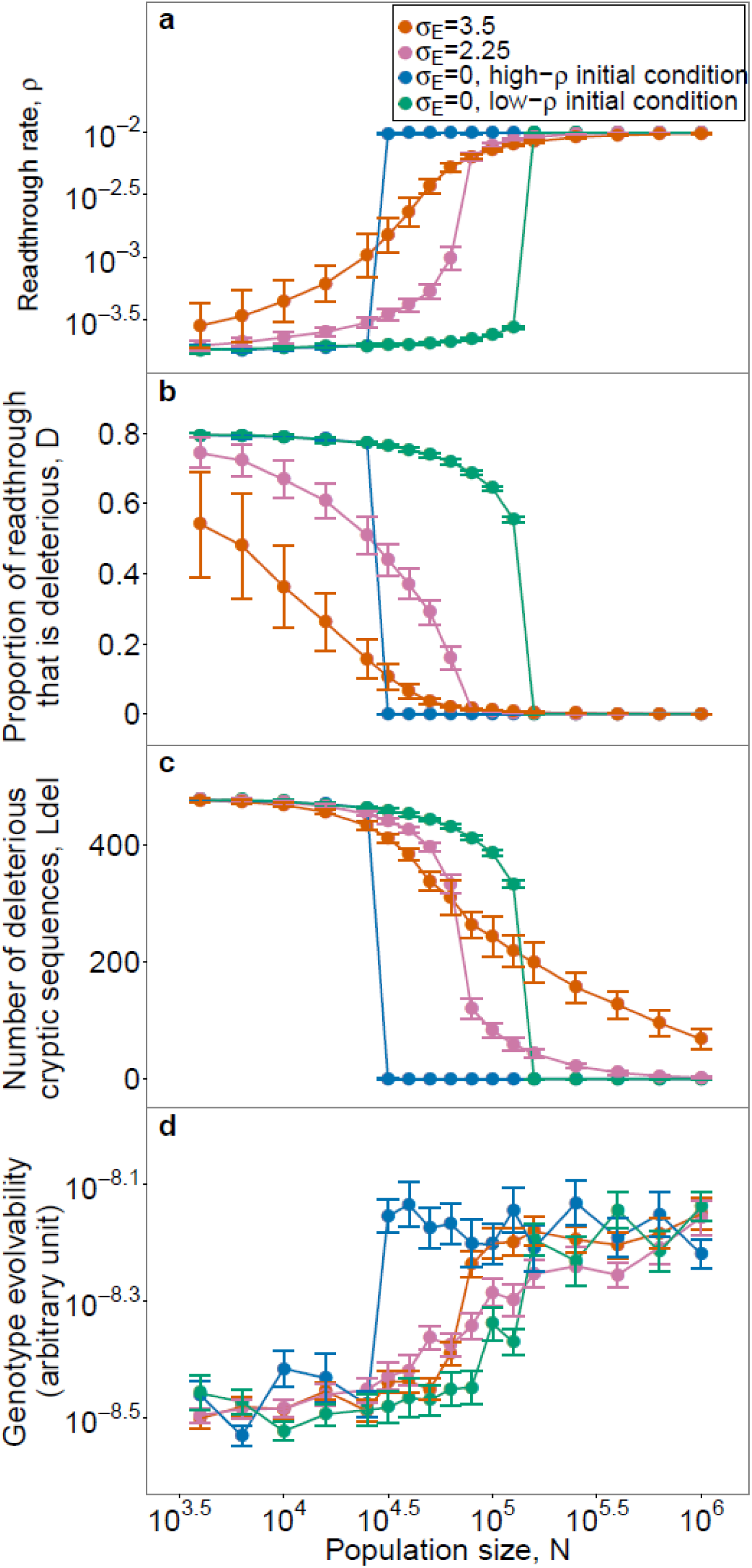
Evolutionary dynamics are bistable in the absence of variation in gene expression (*σ* _*E*_ = 0), but not with variation in gene expression (*σ*_*E*_ = 2.25 and *σ*_*E*_ = 3.5). We calculated the average values of *ρ, D*, and *L*_*del*_ towards the end of the simulations, and then measured the genotype evolvability after changing the optimal trait value (see Methods for details). For each value of *N*, 20 simulations were initialized at high-*ρ* conditions and 15 at low-*ρ* conditions. For *σ*_*E*_ = 2.25 and *σ*_*E*_ = 3.5, simulations from the two initial conditions reached indistinguishable endpoints (Fig. S1), so the results were pooled. The increment in *N* is 10^0.1^ between 10^4.4^ and 10^5.2^ to increase resolution, and is 10^0.2^ elsewhere. At *σ*_*E*_ = 0, *D* is indistinguishable from zero for *N* ≥ 10^5.2^ under high-ρ conditions and for *N* ≥ 10^4.7^ under low-ρ conditions, corresponding to *L*_*del*_ being effectively zero. In contrast, when *σ*_*E*_ = 2.25 or 3.5, because the weakness of selection on low-expression genes prevents *L*_*del*_ from falling all the way to zero, *D* never quite reaches zero either, despite appearing superimposable in **b**. For **a** to **c**, data is shown as mean±SD. For evolvability (**d**), data is shown as mean±SE. For **a** and **d**, these apply to log-transformed values. Evolvability is based on time to fitness recovery; see Fig. S3 for similar results based on time to trait recovery. *L* = 600.

Our results qualitatively reproduce the bistability reported by Rajon and Masel (2011) for the case where there is no expression variation among genes, though the range of values of *N* leading to bistability is smaller than that found in Rajon and Masel (2011) in which a full Wright-Fisher simulation is used. The smaller range of bistability in our model could be caused by the ease with which long-term evolution is captured using an origin-fixation framework, or by other subtle differences between the approaches, e.g. the greater ease of compensatory evolution under Wright-Fisher dynamics than under origin-fixation. We chose origin-fixation mainly to reduce the computational burden, which for our study was increased by the need to track individual loci, in contrast to previous work that needed only to track the number of loci with deleterious cryptic sequence, without distinguishing their identities (Rajon and Masel 2011; Rajon and Masel 2013).

However, bistability vanishes with variation in expression among genes (Fig. 1, *σ*_*E*_ = 2.25 and *σ*_*E*_ = 3.5). To understand why, consider a population initialized at low readthrough rate (*ρ*) and many deleterious cryptic sequences. Because the strength of selection against a deleterious cryptic sequence at locus *i* is proportional to *ρE*_*i*_ (the effect of a locus *i* on *D* in Eq. 3 is proportional to *E*_*i*_), purging works at the most highly expressed loci, even when *ρ* is low. This lowers the proportion *D* of readthrough events that are deleterious, which relaxes selection for high fidelity, leading to an increase in *ρ*. As *ρ* increases, loci with lower *E*_*i*_ become subject to effective purging, which further reduces *D*, which feeds back to increase *ρ* further. Because *E*_*i*_ is log-normally distributed, but contributes linearly to selection via *D*, each round of the feedback loop involves smaller changes than the last. Eventually, the changes are too small for selection on them to overcome mutation bias in favor of deleterious sequences. Similarly, when a population is initialized at high *ρ*, mutational degradation begins at low *E*_*i*_ sites and arrests when selection is strong enough to kick in. The point of balance between mutation bias and selection defines a single intermediate attractor for *σ*_*E*_ ≥ 2.25, instead of the bistable pair of attractors found for uniform *E*_*i*_ (*σ*_*E*_ = 0). For *σ*_*E*_< 2.25, bistability is still found, but for a narrower range of population sizes than in the absence of variation (Fig. S2).

Even though bistability is not found for *σ*_*E*_ = 2.25, there is still a fairly sharp dichotomy, with solutions being either local (high *ρ* and low *L*_*del*_) or global (low *ρ* and high *L*_*del*_), and intermediate solutions found only for a very restrictive range of *N*, following a sigmoidal curve (Fig. 1**a** and 1**c**). Increasing variation in expression among genes blurs the boundary between the local solution and the global solution. Intermediate solutions are found for broader ranges of *N* as expression variance *σ*_*E*_ increases to 3.5. The trend, as expression variance *σ*_*E*_ increases from 0, is to first replace bistability with a limited range of intermediate solutions (*σ*_*E*_ = 2.25), and then for the intermediate solutions to become more prevalent, with extreme local and global solutions becoming less attainable as *σ*_*E*_ > 2.25.

The breakdown of the local solution begins with intermediate values of *L*_*del*_, while the breakdown of the global solution begins with intermediate values of *ρ* and *D* (Fig. 1 **a**-**c**). The breakdown of global solutions involves high-expression loci (Fig. 2), which affect *D* more than *L*_*del*_. In contrast, the breakdown of local solutions involves low-expression loci (Fig. 2), which affect *L*_*del*_ more than *D*. Because *ρ* is better described as co-evolving with *D* than with *L*_*del*_, as explained earlier, intermediate values of *ρ* are seen more in the breakdown of global than local solutions.

**Figure 2:**
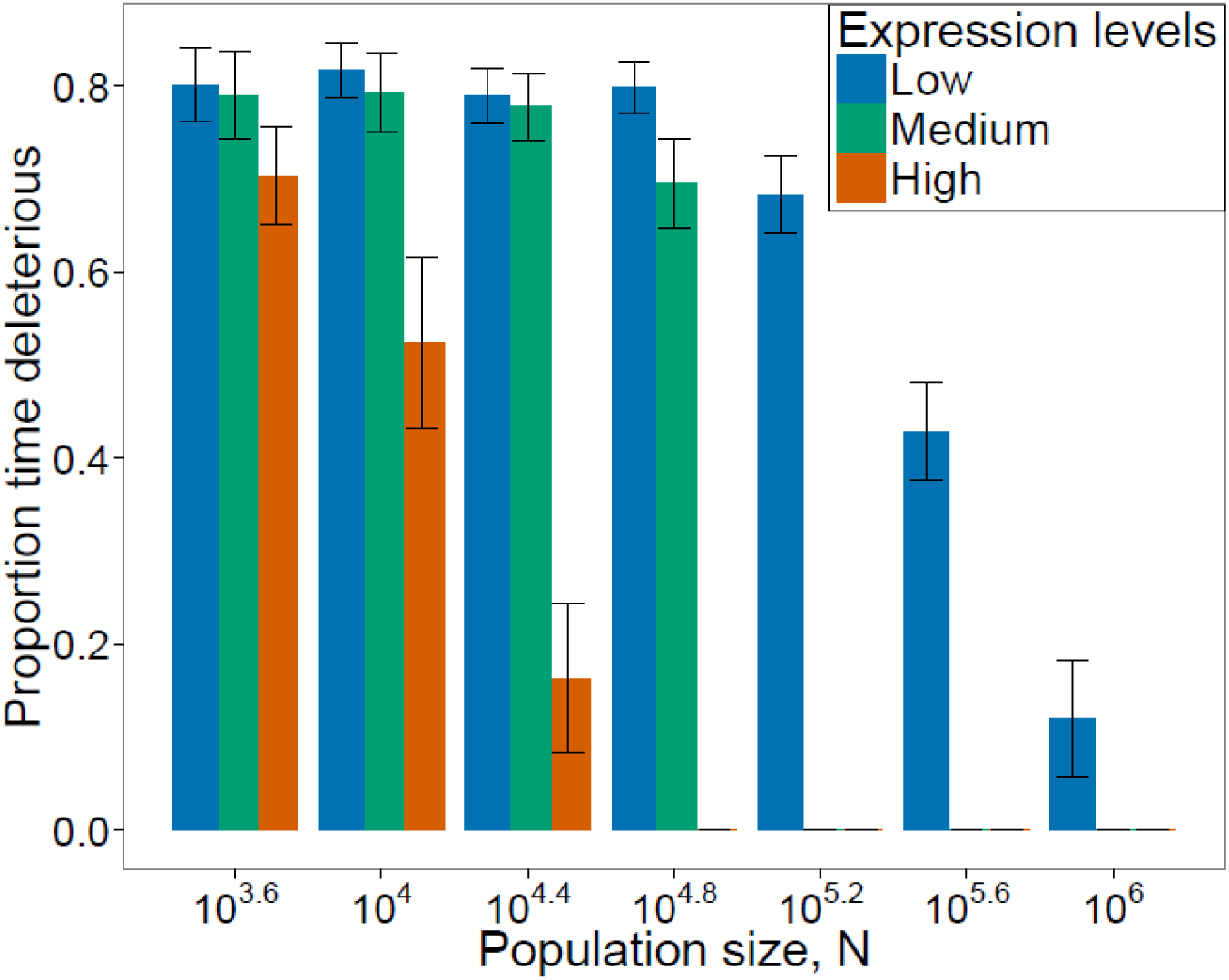
The effectiveness of purging a cryptic sequence of deleterious mutations depends on its expression level. We examined the states of the cryptic sequences of the loci with the 10 highest, the 10 lowest, and the 10 median expression levels among the 600 loci in each of the simulations showed in Fig. 1 (*σ*_*E*_ = 2.25). We counted how often each locus contained a deleterious cryptic sequence among the last 20 timepoints we had collected from that simulation. Bars represent the proportion of time that each of the 10 loci carried a deleterious cryptic sequence, averaged over 20 replicates, and shown as mean ±SD. Simulations were initialized at low-*ρ* conditions.

A primary motivation behind characterizing the two solutions is that the local solution was found to have dramatically higher evolvability than the global solution (Rajon and Masel 2011). We therefore check whether this conclusion still broadly stands in the presence of variance in expression levels. The local solution promotes evolvability by making benign cryptic sequences available for co-option. Differences in evolvability between genotypes should therefore be largely determined by the fraction of quantitative trait loci that carry benign rather than deleterious cryptic sequences. In agreement with this, evolvability inversely mirrors *L*_*del*_, as a function of population size, i.e., evolvability (Fig. 1**d**) resembles *L*_*del*_ (Fig. 1**c**) far more than it resembles *ρ* (Fig. 1**a**) or *D* (Fig. 1**b**).

The distinction between global and location solutions becomes more extreme when the mutation bias toward deleterious rather than benign cryptic sequences is increased from 4:1 ratio to a 99:1 ratio, but persists even when the mutation bias is eliminated in favor of a 1:1 ratio (Fig. 3). In the absence of mutation bias, there is less evolvability to be gained by the local relative to the global solution, since half the quantitative loci are available for co-option regardless (Fig. 3**c**). Nevertheless, a small evolvability advantage to the local solution can still be observed (Fig. 3**d**). In any case, assuming mutation bias toward deleterious options is biologically reasonable, and Fig. 3 shows that results are not sensitive to the quantitative strength of our assumption on this count.

**Figure 3:**
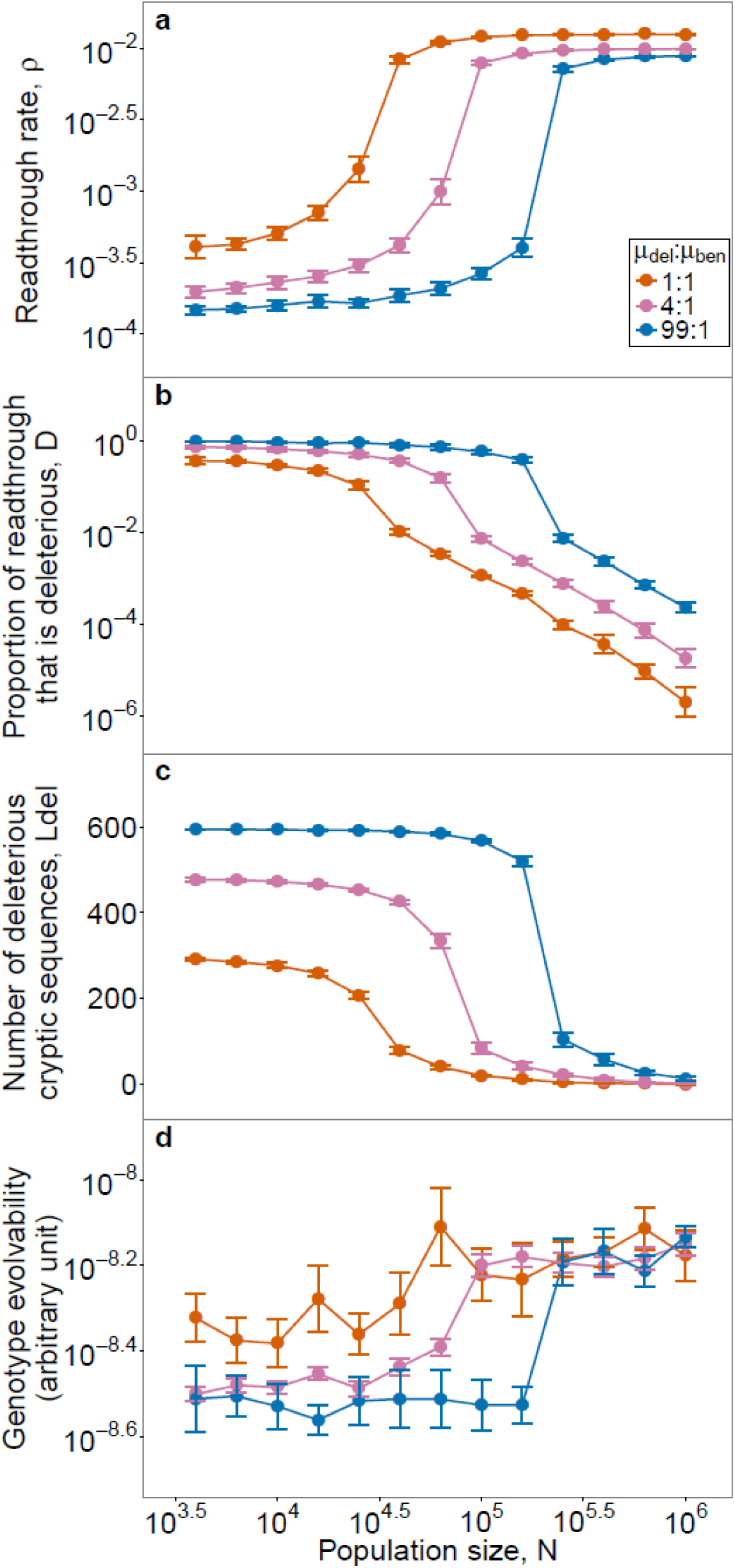
Results become more extreme when the mutation bias in the state of a cryptic sequence is increased from 4:1 ratio to a 99:1 ratio, but do not disappear completely when the mutation bias is eliminated in favor of a 1:1 ratio. The location of the drift barrier shifts as a function of mutation bias, but the dichotomy between local and global solutions (as seen in values of *ρ* and *D*) is not sensitive to relaxing the mutation bias. The advantage of the local solution with respect to evolvability (as seen in **d** and mirrored in *L*_*del*_ (**c**)) is more sensitive to lack of mutation bias, but is still visible even with a 1:1 ratio. To compare results across different mutation biases, we kept the sum of the two mutation rates constant. For the low-*ρ* initial conditions, the number of deleterious cryptic sequences was initialized at the neutral mutational equilibrium of *L*×*μ*_*del*_/(*μ*_*del*_+*μ*_*ben*_) (rounded to the nearest integer). For *μ*_*del*_:*μ*_*ben*_ = 4:1, we reused the results shown in Fig. 1. For the other ratios, five replicates were run for each initial condition, and pooled. For panels **a** to **c**, data is shown as mean±SD. For panel **d**, data is shown as mean±SE. For **a** and **d**, these apply to log-transformed values. *L* = 600 and *σ*_*E*_ = 2.25.

When we also account for mutation bias that tends to increase rather than decrease the error rate *ρ*, our model can explain the previously puzzling observation that the rate of transcriptional errors in small-*N*_*e*_ endosymbiont bacteria *Buchnera* is so much higher than that of *C. elegans,* and almost as high as that of large-*N*_*e*_ *E. coli* (McCandlish and Plotkin 2016; Traverse and Ochman 2016b). In extremely small populations, even the global solution is subject to a drift barrier, making *ρ* higher than its optimal value. For *N* so small such that most *ρ*-increasing mutations pass through the drift barrier, *ρ* can be almost as large as that in large populations (Fig. 4**a**). Despite their high error rates, these extremely small populations also carry heavy loads of deleterious cryptic products (Fig. 4**b** and **c**), consistent with the fact that in *B. aphidicola,* unlike *E. coli,* selection is unable to reduce the fraction of non-synonymous transcriptional errors that are non-synonymous (Traverse and Ochman 2016a). High *ρ* shows the absence of a global solution, while high *D* and *L_del_* show the absence of a local solution; neither solution is found for a sufficiently small population. Similar error rates in large and small populations can also be found, given bias in mutations to *ρ*, when there is no variation in expression levels (Fig. S5).

**Figure 4:**
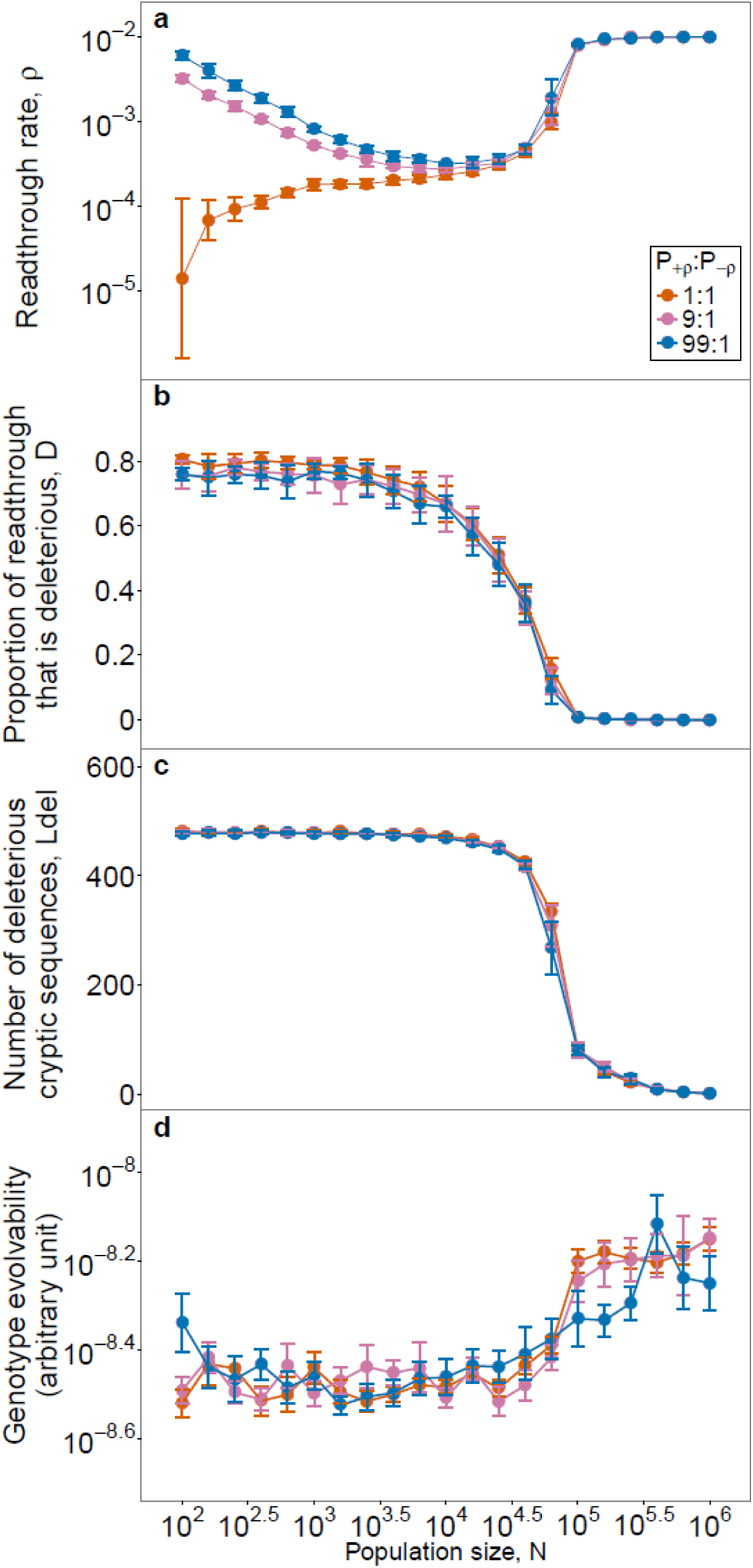
Mutation bias tends to increase *ρ*, such that even the global solution breaks down in sufficiently small populations. *P+p* is the probability that a mutation increases *ρ*, and *P*_-*ρ*_ is the probability of a decrease. Each data point, (except those taken from Fig. 1 with *P*+_*ρ*_:*P*_-*ρ*_ = 1:1 and *N* = 10^3.6^ to *N* = 10^6.0^), is pooled from 5 replicates of high-*ρ* initial conditions and 5 replicates of low-*ρ* initial conditions. Because we assume multiplicative mutational effects to *ρ*, its value converges even for extremely small *N*. I.e., as *ρ* increases, the additive effect size Δ*ρ* of a typical mutation also increases, preventing it from passing through the drift barrier. For **a**, **b**, and **c**, data is shown as mean±SD. For **d**, data is shown as mean±SE. For **a** and **d**, these apply to log-transformed values. *L* = 600 and *σ*_*E*_ = 2.25.

The parameters in our model can be classified into three groups, and the exploration of their values is summarized in Table S1. The first group controls selection coefficients relevant to the global vs. local solution outcome: the variance in expression levels (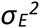), the number of loci (*L*, Fig. S4), the cost of misfolded protein molecules (*c*), and the cost of quality control (*δ*, Fig. S6). The second group controls mutation bias relevant to the global vs. local solution outcome: the frequency with which mutations turn deleterious cryptic sequences benign versus the reverse (*μ*_*ben*_:*μ*_*del*_), whether mutations to *ρ* tend to increase or decrease it (*P*_+*ρ*_:*P*_−*ρ*_), and variance in the magnitude of mutations to *ρ* (*σ*_*ρ*_^2^, Fig. S7). The third group contains all the parameters that control the evolution of quantitative traits encoded by a minority of loci relevant to the evolvability properties. Because our focus in this manuscript is on the evolution of global vs. local solutions, not on the precise details of the relationship between local solutions and evolvability, these parameter values were explored less.

The influence of 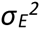 dominates our results. Its effect in eliminating bistability holds, with the one exception that very “cheap” quality control could partially restore bistability (Fig. S6). Otherwise, we found that three parameters – *c*, *δ*, and *μ*_*ben*_:*μ*_*del*_ – are the main determinants of the population size at which the transition between global and local solutions takes place, and of the exact error rate that evolves for global and local solutions (Table S1). The other parameters in the first and second groups have little or no influence on the evolutionary outcomes that we study. In general, parameters in the first group, controlling selection, have stronger effects than the second group, controlling mutation bias.

## DISCUSSION

When genes vary in their expression levels, the dichotomy between the local and global solutions is replaced by a continuous transition. Very large populations still resemble the local solution, although mutations making cryptic sequences deleterious still pass through the drift barrier in the occasional low-expression gene. Very small populations still resemble the global solution, although mutations making cryptic sequences deleterious may still be effectively purged in a few high-expression genes; because their high expression disproportionately affects the burden from misexpression, this relaxes expression for high fidelity, leading to less strict quality control.

In agreement with drift barrier theory, large-*N*_*e*_ *E. coli* exhibits a local solution – a tendency for transcription errors to have synonymous effects – while small-*N*_*e*_ *B. aphidicola* does not (Traverse and Ochman 2016a). While as predicted, the global solution of low transcriptional error rates does not obey the naïve drift barrier expectation of being higher in *B. aphidicola* than in *E. coli* (Traverse and Ochman 2016a), nor are transcription error rates drastically lower in *B. aphidicola* as predicted by previous theory on the interplay between global and local solutions (Rajon and Masel 2011; McCandlish and Plotkin 2016). This significantly lower rate relative to *E. coli* is, however, found in intermediate-*N*_*e*_ *C. elegans.* Where previous work (Rajon and Masel 2011) explained only the relative rates for *E. coli* and *C. elegans,* here we also explain the high error rate of *B. aphidicola* by taking into account a drift barrier on the global solution of low error rates. This drift barrier is significant because of mutation bias towards higher error rates. Small *B. aphidicola* populations have higher error rates than *C. elegans* because it is the best that evolution at low *N*_*e*_ can manage, despite the deleterious consequences; large *E. coli* populations have similarly high error rates because with the worst consequences of error already purged, they don't need to incur the cost that quality control entails.

With small amounts of variation in expression among genes, the range of intermediate values of *N*_*e*_ for which bistability is found shrinks. With more variation, bistability vanishes in favor of a sigmoidal transition between global and local solutions. With still more, the sigmoid is smoothed out, and intermediate solutions are found for most values of *N*_*e*_.

To interpret our results correctly, we must therefore estimate the degree to which genes vary. The results presented here focus on two estimates of the variation in log-expression in yeast, namely standard deviations of 2.25 and 3.5. However, variation among genes in the deleterious consequences of misfolding, in addition to variation in expression levels, might make larger standard deviations a better model of reality, further supporting a continuum of intermediate solutions. In other words, the value of *c* in Eq. 3 may vary among genes. Note that apart from the second-order *ρ*^2^ term, the cost of a deleterious misfolded protein *i* depends only on the product of *c*_*i*_ and expression level *E*_*i*_. Given log-normal distributions of *c*_*i*_ and expression level *E*_*i*_, the variance of the log-product is equal to the sum of the two log-variances, so we can transform this scenario into one where *c* is constant, and 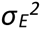 is equal to this sum. This can be done because changing *c*_*i*_ and *E*_*i*_ only affects *w*_*misfolding*_ and not other factors such as the magnitude of a locus's influence on the quantitative trait. In other words, adding variation to *c* is almost equivalent to increasing the variance in expression levels.

The values of *μ*_*del*_ and *μ*_*ben*_ may also vary among genes. Drift barrier effects operate via the effect of population size on the fate of deleterious not beneficial mutations – if purging is efficient, then the beneficial mutation rate does not matter, because a single beneficial mutation is enough. We therefore focus on *μ*_*del*_. The inclusion of a benign-to-deleterious mutation *M*_*i*_ at locus *i* depends on the product of *μ*_*del*_ at locus *i* and *M*_*i*_'*s* probability of fixation. It seems likely that variation among genes in the probability that a deleterious cryptic sequence becomes fixed will swamp variation in the deleterious mutation rate – variation in expression levels cause the former to vary over orders of magnitude. Note that as for the case of variation in *c*, it is possible to construct a manipulation of *E*_*i*_ that has the same effect on the relevant product, via the probability of fixation, as would occur given a change in *μ*_*del*_. While this case is less neat than for the product *c*_*i*_*E*_*i*_, it illustrates that a model of variation in expression levels can reflect, to some extent, the effect of variation in *μ*_*del*_.

Our model makes three critical assumptions, which must be understood for the results to be interpreted appropriately. First, a “locus” in our model consists of one regular and one cryptic sequence. The primary example that we used to parameterize the simulations posits an entire protein-coding gene as the regular sequence, and the extended polypeptide resulting from stop codon readthrough as the cryptic alternative. In the example of transcriptional errors, a locus is a single codon, with its corresponding amino acid being the regular sequence, and the most common consequence of a transcriptional error as the cryptic. The case of one regular sequence and many alternative cryptic ones has not been modeled. Similarly, proteins may each have a regular fold or binding partner, and our model considers the contrast between this state and a single cryptic alternative.

Second, we assume that the rate of gene expression errors is set globally, across all loci. In reality, individual context may also affect the error rate, giving error rates a local solution aspect as well. A model of three rather than two interacting solutions – global error rates, local error rates, and local robustness to the consequences of error – remains for future work. Perhaps highly expressed genes will have both more benign cryptic sequences and lower rates of error, or perhaps the evolution of one kind of local solution will alleviate the need for another. Testing this empirically requires data on site-specific error rates and on a credible marker for the benign status of members of an identifiable class of cryptic sequences. Such tools are now becoming available, and indeed we recently found a positive correlation between a large number of readthrough errors at a particular stop codon and the benign status of the readthrough translation product (Kosinski et al., manuscript in preparation). We also reanalyzed the data of Traverse and Ochman (2016a) to find that highly expressed transcripts have lower transcriptional error rates (unpublished result).

Finally, we assume that the consequences of errors have a bimodal distribution: either highly deleterious or largely benign, but rarely in between. In other words, we assume that a basic phenomenon in biology is that changes tend to either break something, or to tinker with it. There are a variety of lines of evidence supporting this intuitively reasonable assumption (Fudala and Korona 2009; Wylie and Shakhnovich 2011).

## ACKNOWLEDGEMENTS

This work was supported by the John Templeton Foundation [grant number 39667]. DJP was also funded by the Undergraduate Biology Research Program at the University of Arizona. We thank Lilach Hadany, Yoav Ram, and other members of the Hadany group for helpful discussions that prompted us to explore variation in expression. We thank Paul Nelson for developing the idea of local error rates as an expansion of our model, Ben Wilson for help with R, and Etienne Rajon, Yoav Ram, Tobias Warnecke, and one anonymous reviewer for helpful comments on the manuscript. We also thank Charles Traverse and Howard Ochman for sharing their data on transcriptional errors. An allocation of computer time from the UA Research Computing High Performance Computing (HPC) and High Throughput Computing (HTC) at the University of Arizona is gratefully acknowledged.

## Supplemental material for the manuscript “Drift barriers to quality control when genes are expressed at different levels”

Xiong K, McEntee JP, Porfirio DJ, Masel J

### Implementation of origin-fixation simulations

Origin-fixation models are often implemented via a crude rejection algorithm; large numbers of mutations are simulated, and each is accepted as a successful fixation event if and only if a random number sample from the uniform [0, 1] distribution falls below its (fairly low) fixation probability. For large *N*, this method is computationally slow when significant numbers of nearly neutral mutations must be sampled before one fixes with probability ∼1/*N*. Given that our model posits only a relatively small range of possible mutations, we instead sampled only mutations that go on to become fixed, by sampling according to the relative values of “fixation flux”, proportional to mutation rate × fixation probability for each of our six categories of mutation. In other words, we used a form of the Gillespie (1977) algorithm.

In a haploid population of size *N*, the probability of fixation of a new mutant into a resident population is given by

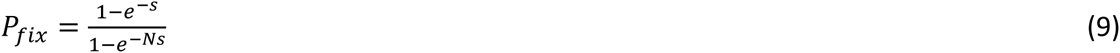

where *s* = *w*_*mutant*_/*w*_*resident*_−1. It is then straightforward to calculate fixation flux values for all possible switches between benign and deleterious states:

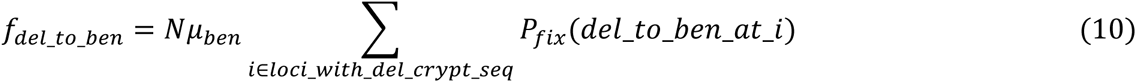

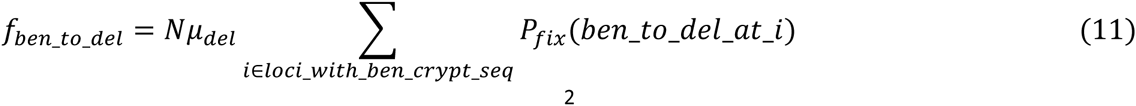

Matters are slightly more complicated for quantitative mutations to *α*, *β* and *ρ*, because we must integrate the fixation flux over all possible sizes (Δ*α*_*k*_, Δ*β*_*k*_, and Δlog_10_*ρ*) for a mutation at a given locus, prior to summing across loci to arrive at the fixation flux for an entire mutational category:

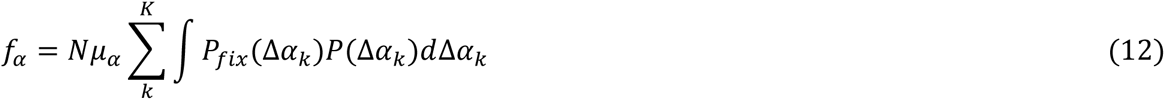

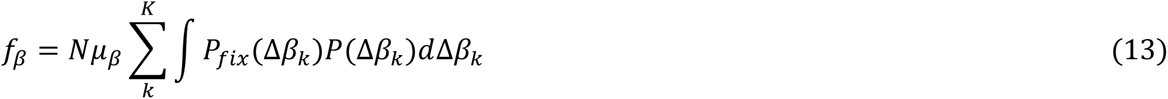

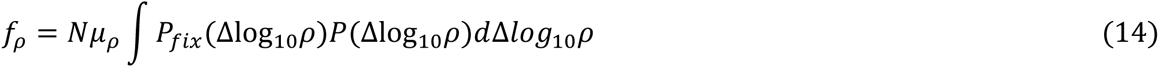

where *P*(Δ*α*_*k*_), *P*(Δ*β*_*k*_), and *P*(Δlog_10*p*_) are the probability densities for the magnitude of a given kind of mutation.

We use the quadrature method to calculate the integral over these possibilities, using a grid of 2000, limited for Δ*α*_*k*_ to the interval [–*α*_*k*_/a-5*σ*_*m*_/*K*, –*α*_*k*_/a+5*σ*_*m*_/*K*], for Δ*β*_*k*_ to the interval [–*β*_*k*_/*a*-5*σ*_*m*_/*K*, –*β*_*k*_/a+5*σ*_*m*_/*K*], and for Δlog_10*ρ*_, to the interval [−10*σ*_*ρ*_, min(10*σ*_*ρ*_, −log_10*ρ*_)]. In the latter case, the number of grid intervals is reduced proportional to any truncation of the interval at −log_10_*ρ*.

For mutational co-options of benign cryptic sequences, the effect of replacing the value of *α*_*k*_ with that of *α*_*k*_+*β*_*k*_ is fixed, but there is also a stochastic range of effects of initializing a new *β*_*k*_ and a new *B*_*k*_(Eq. 15). Let *P*(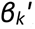) be the probability density of a new *β*_*k*_given by Normal(0, *V*(*a*, *K*, *σ*_*m*_)), and *P*(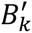 = 1) = 1 – *P*(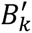 0) be the probability that a new *B*_*k*_ equals to 1, and hence the new *β*_*k*_ affects the trait value. The fixation flux associated with cooption mutations we obtained numerically by integration over the range [-5*σ*_*m*_/*K*, 5*σ*_*m*_/*K*]:

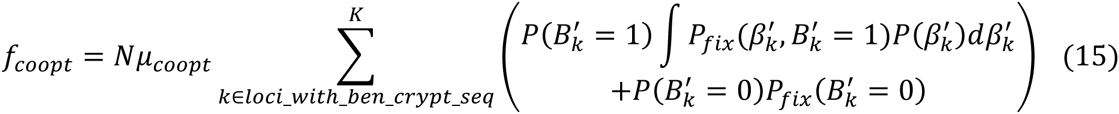

The expected waiting time before the current genotype is replaced by another is

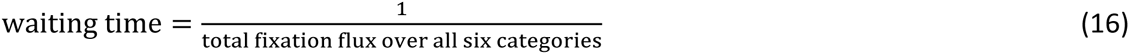

A standard Gillespie (1977) algorithm would calculate the realized waiting time as a random number drawn from an exponential distribution with this mean. Since we are only interested in the outcome of evolution, and not the variation in its timecourse, we used the expected waiting time instead, decreasing our computation time. The waiting time can be interpreted as the time it takes for a mutation destined for fixation to appear, neglecting the time taken during the process of fixation itself. Using this interpretation, we specify waiting times in terms of numbers of generations, based on our assumptions about absolute mutation rates.

We assign the identity of the next fixation event among the six categories according to probabilities proportional to their relative fixation fluxes, then we assign the identity within the category. For switches between benign and deleterious states, allocating a fixation event within a category according to the relative values of fixation fluxes is straightforward. For mutations to *ρ*, *α*, and *β*, and mutational co-option, we relax the granularity and cutoff assumptions of the grid-integration method when choosing a mutation within the category. Instead, we sample a mutational value of Δlog_10_*ρ* from Normal(*ρ*_*bias*_, 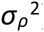). We reject and resample Δlog_10_*ρ* if Δlog_10_*ρ* ≥ – log_10_*ρ*. Otherwise, we accept vs. reject-resample according to the fixation probability of that exact mutation, by comparing this probability to a random number uniformly distributed at [0, 1.1×the maximum fixation probability across the grid points previously calculated for Δlog_10_*ρ* during our grid calculation].For Δ*α* (or Δ*β*), the procedure is conceptually similar but has a more complicated implementation. We first sample from Normal(0, (*σ*_*m*_ /*K*)^2^). We then add the random number to each of the values of –*α*_*k*_/*a*, and calculate the sum of corresponding fixation probabilities across all loci *k*. We accept vs. reject-resample the mutation by comparing this sum to a random sample from a uniform distribution at [0, 1.1×the maximum corresponding fixation probability sum calculated during our grid calculation]. If the mutation is accepted, we allocate it to a locus *k* with probability proportional to their relative fixation probabilities. For mutational co-option of a benign cryptic sequence, the main effect is to replace *α*_*k*_ with *α*_*k*_+*β*_*k*_, but there are also subtler effects arising from the reinitialization of the new cryptic sequence. Any of the *k* loci for which *B* = 1 are eligible for co-option, the new value of *B* may be either 0 to 1, and the new *β*_*k*_ may take a range of values. Each combination of *k* and new *B* has its own fitness flux, and the first choice is among these {*k*, *B*} pairs. Next we sample *β*_*k*_ from Normal(0, (*σ*_*m*_/*K*)^2^); for a new *B* equal to 0 we always accept the result, and for new *B* equal to 1, we accept vs. reject-resample *β*_*k*_ by comparing its probability of fixation to a random sample from a uniform distribution at [0, 1.1×the maximum corresponding fixation probability sum calculated during our grid calculation].

**Figure S1:**
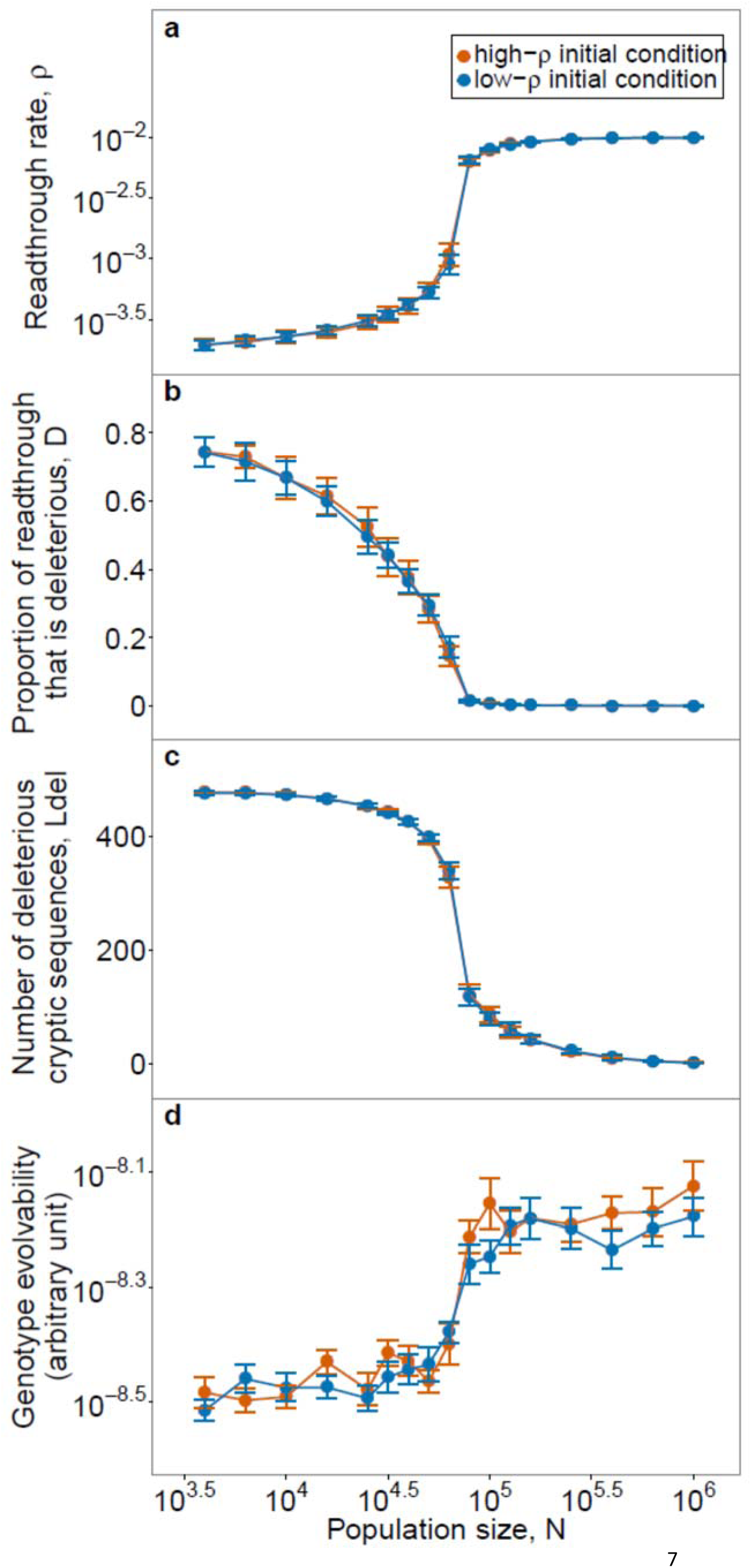
At *σ*_*E*_ = 2.25, the final state of the evolutionary simulation does not depend on the initial conditions. The data shown here is the same as that shown pooled in Fig. 1.

**Figure S2:**
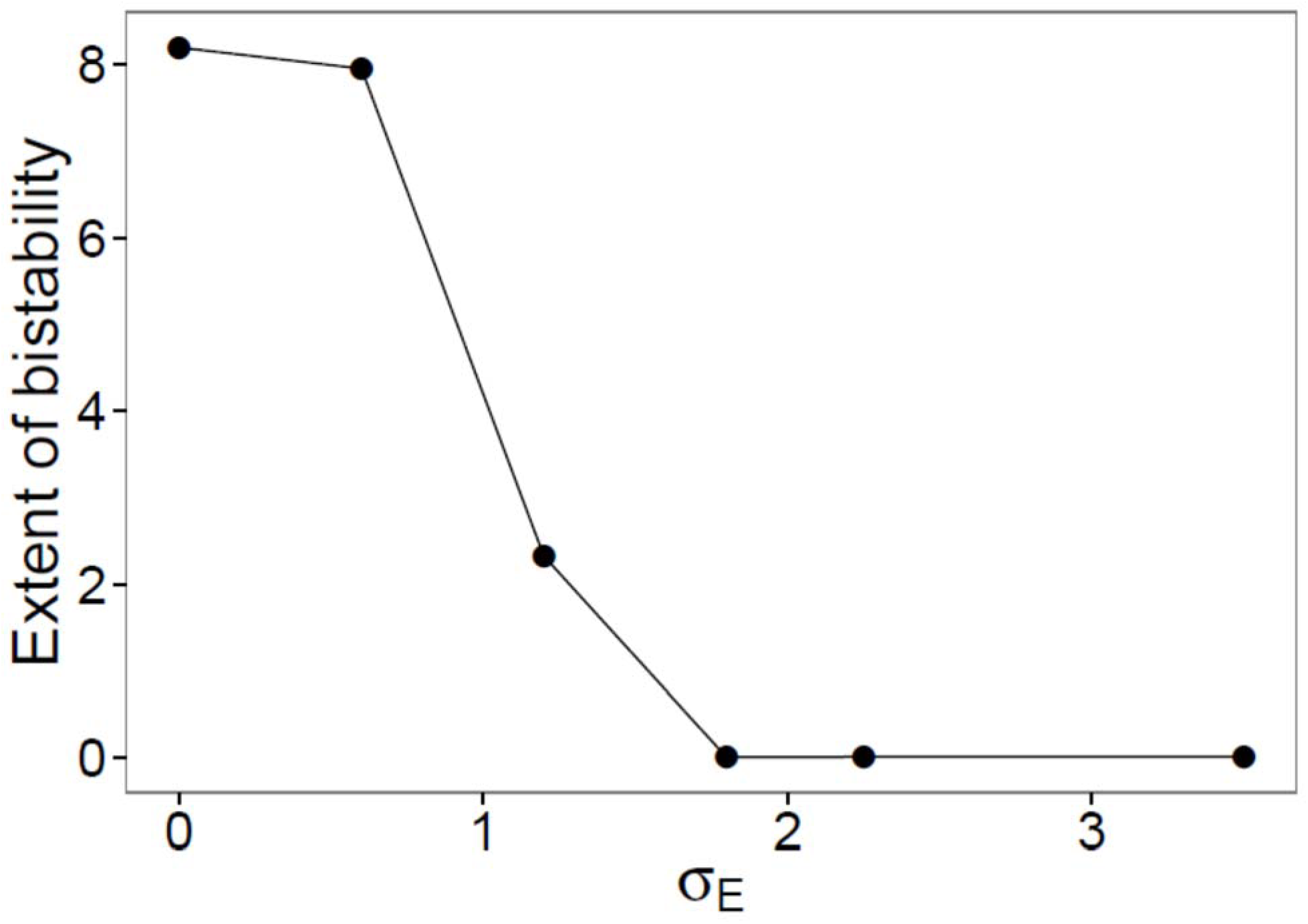
The range of population sizes that exhibit significant bistability drops dramatically even for *σ*_*E*_ < 2.25. We used average values of *ρ* towards the end of the simulations as a measure of the solution found by each replicate. For each initial condition, we averaged over five replicates (except for *σ*_*E*_ = 0, 2.25, and 3.5, where we reused the 20 replicates of Fig. 1), and over each of the values of *N* between 10^3.6^ to 10^6^, with an increment of 10^0.2^. The extent of bistability was assessed as 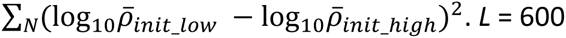.

**Figure S3:**
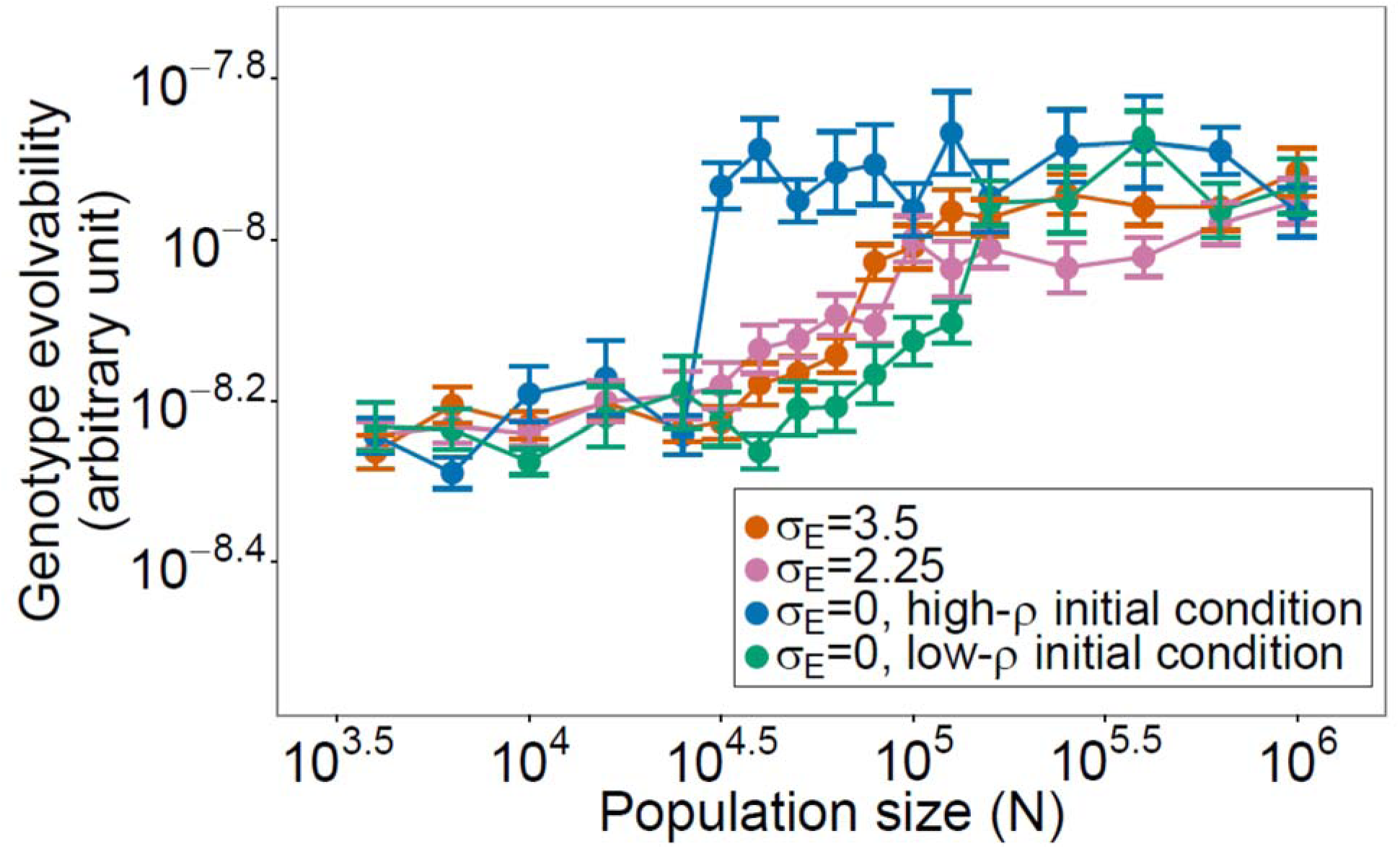
The time taken for the trait to approach the new value of *x*_*opt*_ behaves similarly to the recovery time of fitness shown in Fig. 1**d**. The same simulations were used as in Fig. 1. At *σ*_*E*_ = 2.25 and *σ*_*E*_ = 3.5, we pooled the results from high-*ρ* and low-*ρ* conditions. Evolvability is shown as mean±SE of the log-transformed values. *L* = 600.

**Figure S4:**
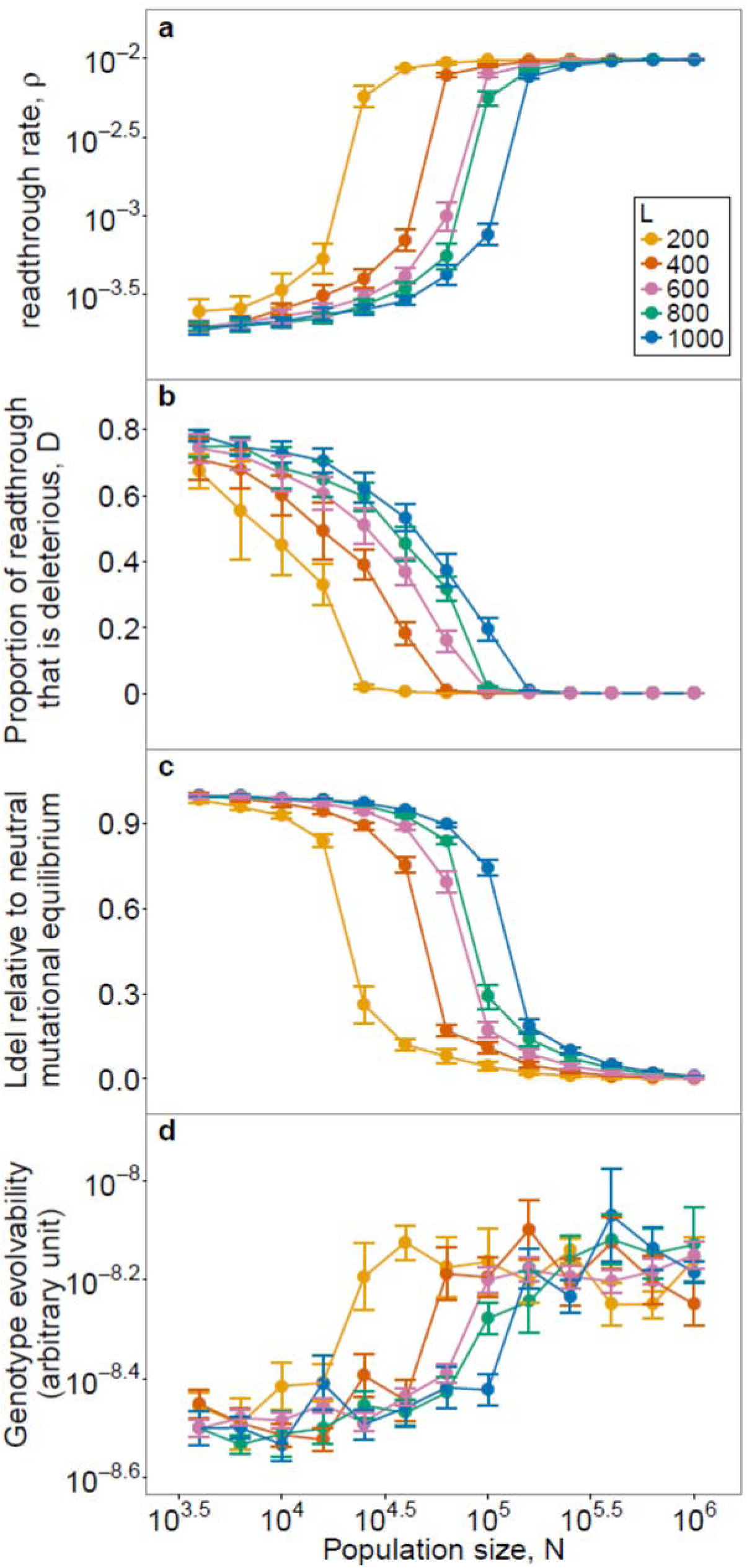
Changing the number of loci does not qualitatively change our results. Quantitatively, fewer loci favor more local solutions. Changing *L* alters the average contribution of each locus to *D*. This alters the average strength of selection on each locus, independent of population size. Therefore, the same solutions, characterized by the values of *ρ* and *D*, are “shifted” to small values of *N* as *L* decreases. While *L* changed, we held the number of quantitative trait loci constant at 50. For *L* = 600, we reused the results shown in Fig. 1. For other values of *L,* five replicates were run for each of the two initial conditions. We pooled results from both initial conditions across all values of *L*. We normalized *L*_*del*_to the neutral mutational equilibrium of *L*×*μ*_*del*_/(*μ*_*del*_+*μ*_*ben*_). For panels **a** to **c**, data is shown as mean±SD. For **d**, data is shown as mean±SE. For **a** and **d**, these apply to log-transformed values. *σ*_*E*_ = 2.25.

**Figure S5:**
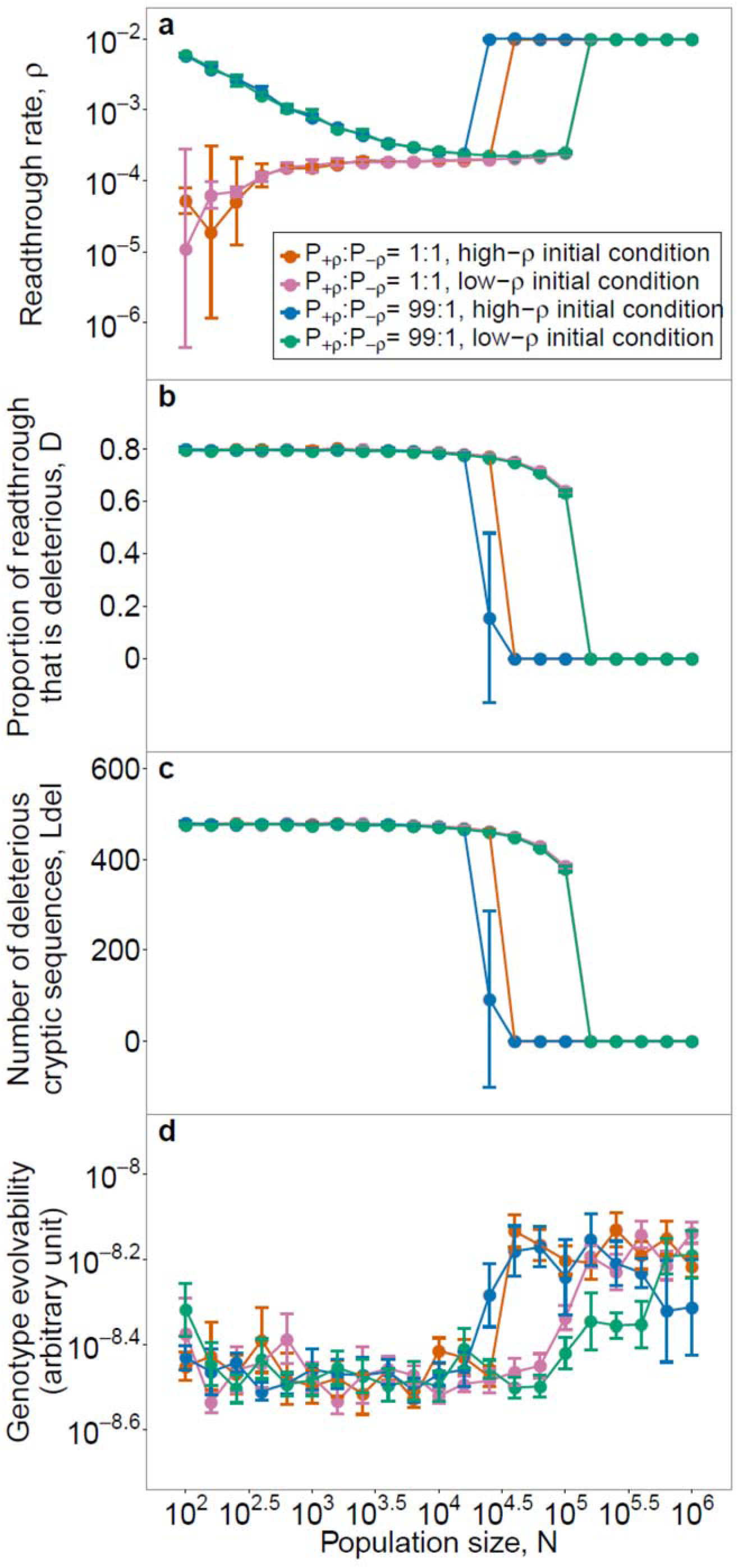
Fig. 4 results (that the global solution breaks down in sufficiently small populations) remain true in the absence of variation of expression levels. Data points between *N* = 10^3.6^ to *N* = 10^6.0^ and *P*_+*ρ*_:*P*_−*ρ*_=1:1, are reused from Fig. 1; for the others, we performed 5 replicates for each condition. For panels **a** to **c**, data is shown as mean±SD. For **d**, data is shown as mean±SE. For **a** and **d**, these apply to log-transformed values. *L* = 600.

**Figure S6:**
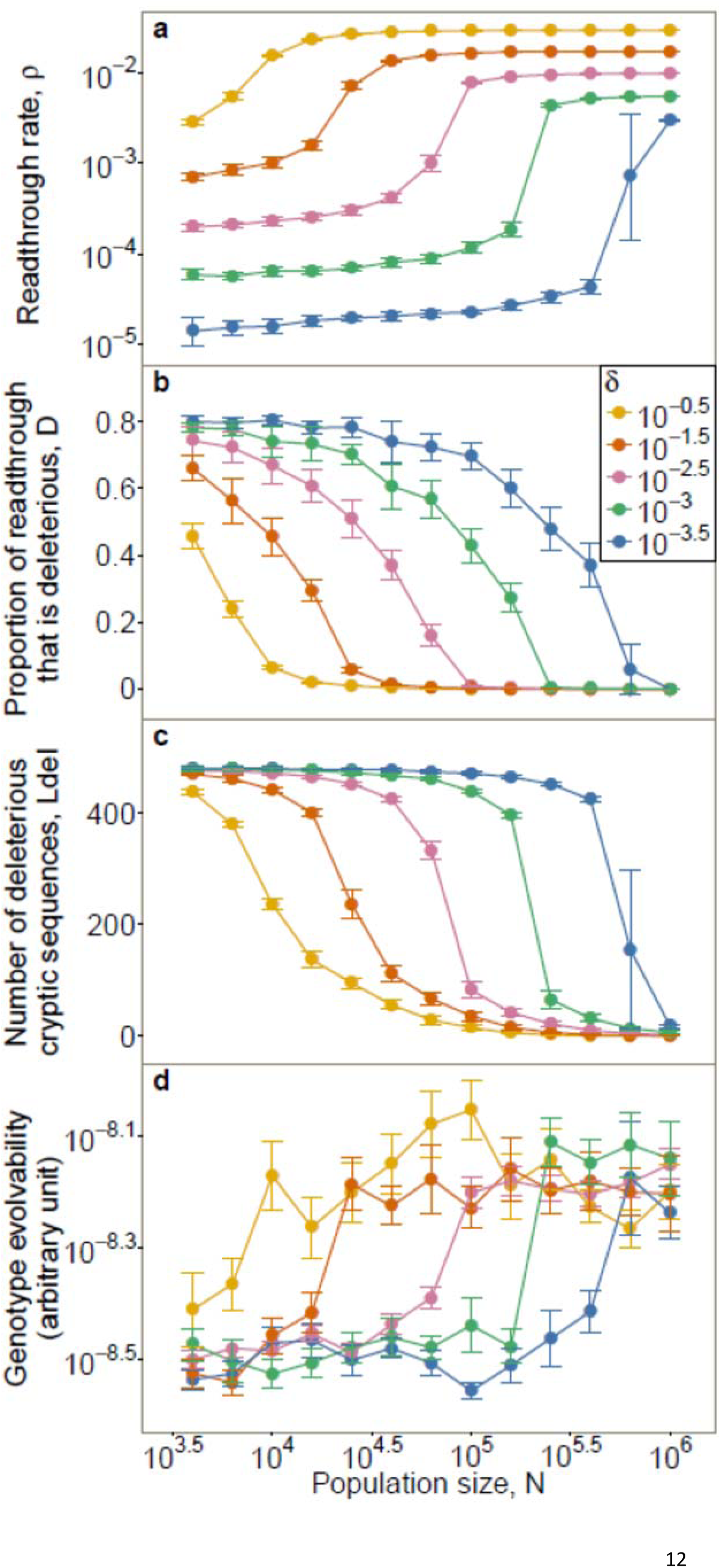
Increasing the cost of quality control *δ* expands global solutions to smaller populations and reduces the differences in error rates as a function of population size. For *δ* = 10^−2.5^, we reused the data from Fig. 1; for each of the other values of *δ*, we ran 5 replicates from the high-*ρ* initial condition and 5 from the low-*ρ* initial condition. Each data point represents the pooled results from the two initial conditions. For panels **a** to **c**, data is shown as mean±SD. For **d**, data is based on time to fitness recovery and is shown as mean±SE. For **a** and **d**, the mean, SD and SE are calculated on log-transformed values. The large error bars at *N* = 10^5.8^ under *δ* = 10^3.5^ across all panels are due to different initial conditions, which is a sign of bistability. *L* = 600, *σ*_*E*_ = 2.25.

**Figure S7:**
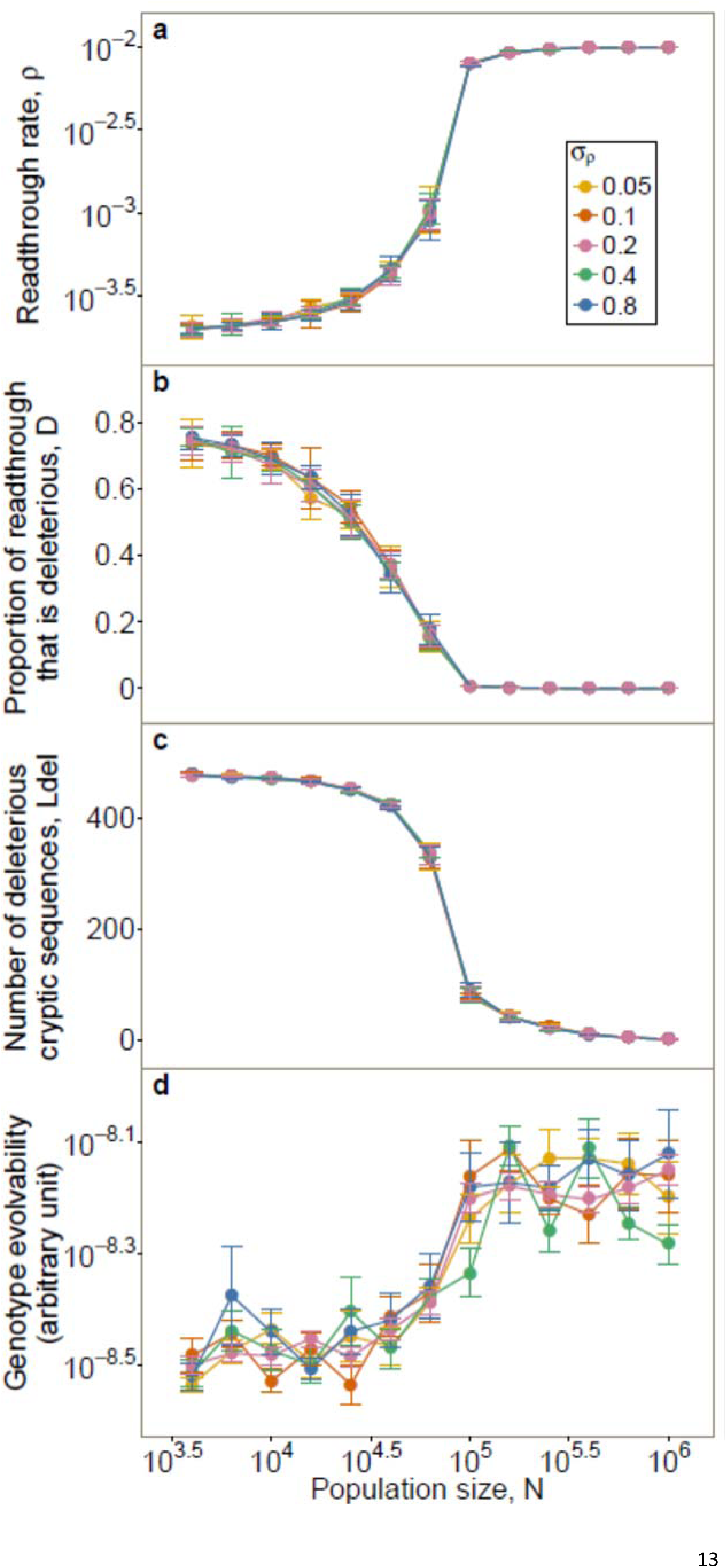
The variance in the magnitude of mutations to *ρ* does not affect a population's solution to error or evolvability. For *σ*_*ρ*_ = 0.2, we reused the data from Fig. 1; for each of the other values of *σ*_*ρ*_, we ran 5 replicates from each of the two initial conditions. We pooled results from the two initial conditions for each data point. For panels **a** to **c**, data is shown as mean±SD. For **d**, data is based on time to fitness recovery and is shown as mean±SE. For **a** and **d**, these apply to log-transformed values. *L* = 600, *σ*_*E*_ = 2.25.

**Table S1.**
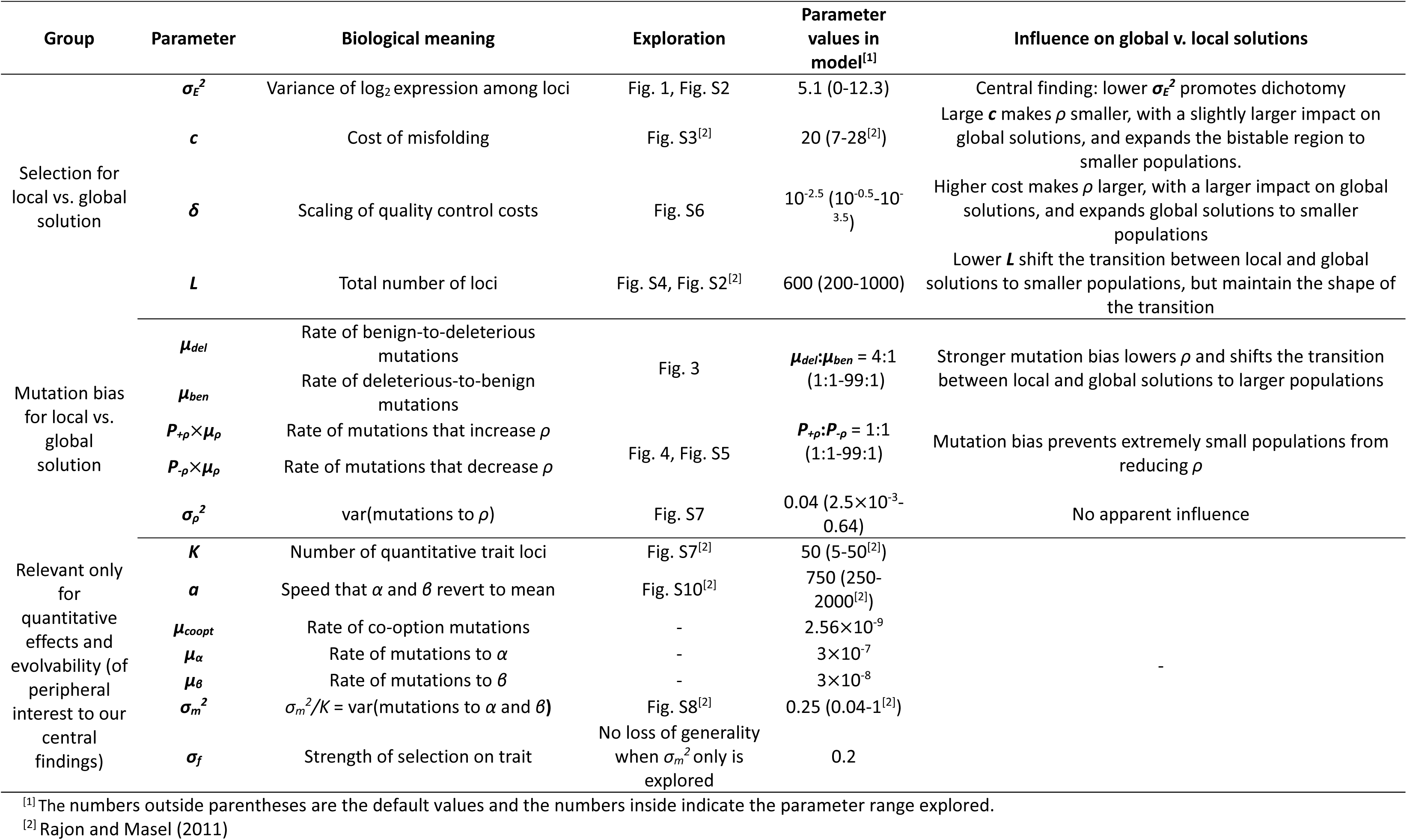
Summary of model parameters

